# Decrypting the complex phenotyping traits of plants by machine learning

**DOI:** 10.1101/2024.11.14.623623

**Authors:** Jan Zdrazil, Lingping Kong, Pavel Klimeš, Francisco Ignacio Jasso-Robles, Iñigo Saiz-Fernández, Firat Güder, Lukaš Spíchal, Václav Snášel, Nuria De Diego

**Affiliations:** Faculty of Electrical Engineering and Computer Science, VSB-Technical University of Ostrava, Ostrava, Czech Republic.; Czech Advanced Technology and Research Institute (CATRIN), Palacký University Olomouc, Slechtitelů 27, 77900, Olomouc, Czech Republic.; Department of Bioengineering, Imperial College London, London SW7 2AZ, UK.

## Abstract

Phenotypes, defining an organism’s behaviour and physical attributes, arise from the complex, dynamic interplay of genetics, development, and environment, whose interactions make it enormously challenging to forecast future phenotypic traits of a plant at a given moment. This work reports AMULET, a modular approach that uses imaging-based high-throughput phenotyping and machine learning to predict morphological and physiological plant traits hours to days before they are visible. AMULET streamlines the phenotyping process by integrating plant detection, prediction, segmentation, and data analysis, enhancing workflow efficiency and reducing time. The machine learning models used data from over 30,000 plants, using the *Arabidopsis thaliana-Pseudomonas syringae* pathosystem. AMULET also demonstrated its adaptability by accurately detecting and predicting phenotypes of *in vitro* potato plants after minimal fine-tuning with a small dataset. The general approach implemented through AMULET streamlines phenotyping and will improve breeding programs and agricultural management by enabling pre-emptive interventions optimising plant health and productivity.

## 1. Introduction

Although the genome codes an organism’s potential trait, phenotypes are the real endpoint characteristics that determine an organism’s behaviour and physical attributes at any given time^1,2^. Unlike the well-defined, indivisible nature of genes, phenotypes are infinitely variable, influenced by genetics, developmental stage, and environment. This complexity is magnified by the enormous genetic diversity in the biological world, further expanded by the new genotypes obtained by the traditional and modern human breeding practices^2^.

Humans rely on plants directly, as they constitute 80% of the global diet, 29% of the textile fibres^3^ and 11% of building material^4^ or indirectly, as a source of energy and oxygen and through the consumption of plant-eating animals as a source of food, (https://www.fao.org/plant-health-2020/home/en/). Breeding to modify specific phenotypic traits is challenging due to the diverse environmental conditions of a crop’s life cycle, affecting everything from germination to yield^5^. Consequently, selecting desirable phenotypes is a numbers game hindered by a fundamental lack of biological understanding, making it essential to test many genotypes under various conditions to find the ones that produce the desired phenotypes efficiently.

High-Throughput Phenotyping (HTP) is an advanced method combining imaging, robotics, and sensor networks to collect big datasets from large plant populations^6,7^. An advanced HTP setup can analyse over 25,000 plants in one run^8^, facilitating significant breakthroughs in identifying and characterising biostimulants^9,10^, eco-friendly agrochemicals^11^, and genotypes with improved resilience^12^. Combined with other -omics, HTP also enables an understanding of plant responses to multifactorial stress conditions^13^.

*Arabidopsis thaliana*, a Brassicaceae family member, is the most used model organism in plant research worldwide because of its well-characterised genome, ease of cultivation and manipulation, short life cycle, and prolific seed production plant^14^. Despite the capabilities of miniaturising HTP technologies using Arabidopsis as a model plant, the challenge remains in segmenting these tiny plants (2-4 mm diameter of four-day-old seedling) at earlier developmental stages, and analysing and interpreting these complex datasets is time-consuming and demands advanced statistical and computational techniques beyond traditional methods. For more efficient use of the big data obtained from HTP experiments, tasks such as plant detection, segmentation, and data analysis are crucial for identifying the most relevant characteristics related to plant resilience.

The integration of machine learning (ML) with big datasets from HTP is revolutionising the analysis of complex data from plants grown under different conditions^15,16^. Imaging and ML technologies have proven effective in detecting and classifying plant diseases^15,17^ and response to specific abiotic stressors^18–20^. Regardless of the sophistication of the current approaches offering identification of desirable traits related to plant resistance^21^, most works focus on a single task such as classifying genotypes or diseases, plant segmentation, stomata analysis^21–24^, etc. In other words, current approaches lack versatility as they are tailored to specific procedures or plant species. General-purpose models that can easily be fine-tuned across various experimental pipelines to enhance the versatility of analysis are, therefore, the "holy grail" of plant phenotyping. Moreover, many studies combining ML with plant phenotyping rely on sophisticated and costly non-invasive sensors (fluorescence, thermal, or hyperspectral cameras^19,20,25^), which may not be accessible to all researchers and growers. Furthermore, there is a notable gap in predictive analytics, offering models with substantial potential for forecasting plant changes before they become apparent.

In this study, we introduce **AMULET-A**daptable and **MU**lti-task machine **L**earning for **E**ndless phenotyping **T**rait prediction- the next generation in HTP technology **(Figure 1),** which combines simple RGB imaging with AI-driven models for predicting morphological and physiological traits hours to days before their visible manifestation. This modular pipeline integrates plant detection, segmentation, prediction, and data analysis, making it readily adaptable for different plant species to monitor and predict plant growth and stress response. Fully compliant with REFORMS (REcommendations FOR Machine-learning-based Science) guidelines^26^, AMULET transcends traditional computer vision tools by using longitudinally acquired plant imagery for a comprehensive range of tasks within a unified pipeline (Figure 1), a significant advancement in plant phenotyping, enhancing both its efficiency and impact, thereby supporting the development of more resilient plants and sustainable, productive agricultural practices.

**Figure 1.**
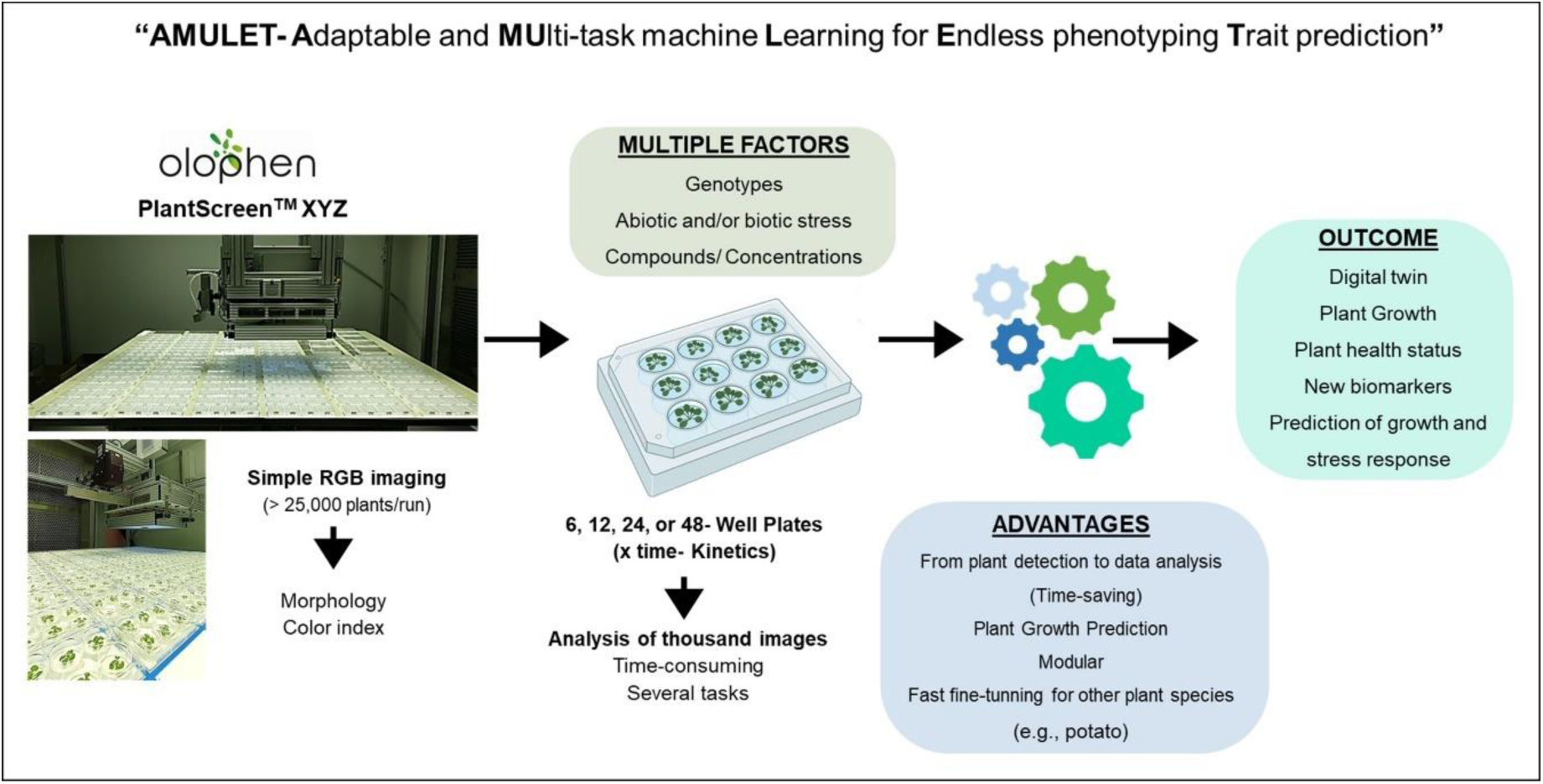
**AMULET**- **A**daptable and **MU**lti-task machine **L**earning for **E**ndless phenotyping **T**rait prediction-pipeline involves different modules, starting with plant imaging using an RGB camera and then selecting different machine learning models for plant detection, prediction, segmentation, and data analysis. The advantages and outcomes are indicated.

## 2. Results and discussion

### 2.1 Workflow of AMULET

The AMULET workflow **(Figure 2 and Supplementary Figure S1)** starts with the automated capturing of high-resolution RGB images of *Arabidopsis thaliana* plants grown *in vitro* into varied well-plates at the Olophen system (http://www.plant-phenotyping.org/db_infrastructure#/tool/57), a plant phenotyping platform equipped with PlantScreen^TM^ XYZ and Compact systems (Photo System Instruments, PSI, Czech Republic). The HTP screening method used the high-resolution top-view RGB camera installed on the PlantScreenXYZ^TM^ to track Arabidopsis growth meticulously over 7 to 10 days^8,9^. This process involves imaging the plants at various developmental stages grown on various multi-well plates to generate a comprehensive training dataset. The main focus was the study of plant-microorganism interaction using the *Arabidopsis thaliana-Pseudomonas syringae* pathosystem as a model and an example of the AMULET power.

**Figure 2.**
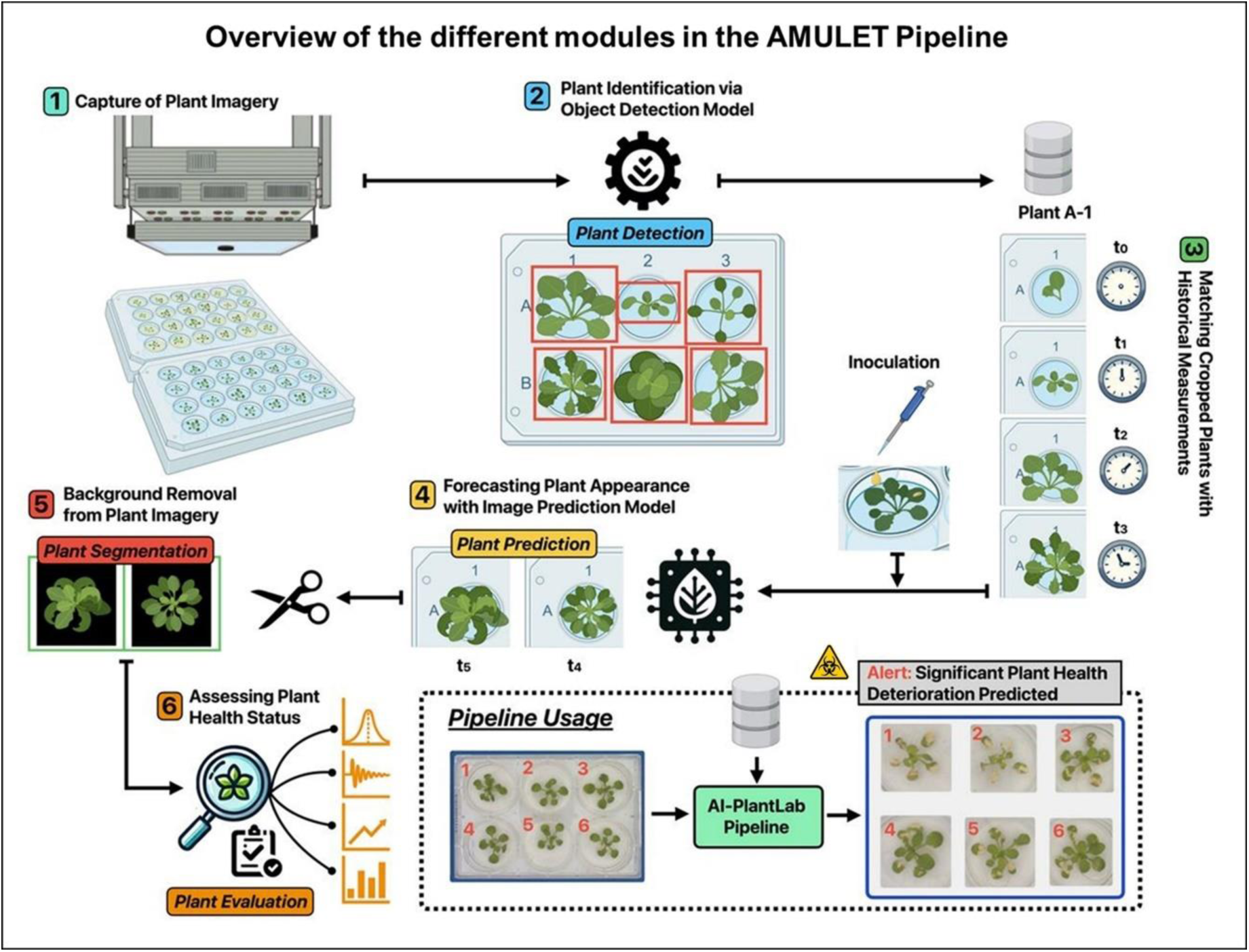
The illustration presents the six stages of the AMULET workflow: (1) Capture of Plant Imagery: High-resolution RGB images of *Arabidopsis thaliana* plants are obtained. (2) Plant Identification via Object Detection Model: Plants are localised and framed within the images. (3) Matching Cropped Plants with Historical Measurements: Cropped plants are paired with historical data, creating a time series. (4) Forecasting Plant Appearance with Image Prediction Model: Future plant growth is predicted based on historical images. (5) Background Removal from Plant Imagery: Segmentation isolates the plant rosette from the background. (6) Assessing Plant Health Status: Plant health is evaluated using a model trained on morphological, physiological, and color descriptors. The pipeline usage is showcased in the bottom right of the image, depicting the input image and the resulting predictions.

AMULET integrates a complex workflow of six stages, including plant imaging, object *detection*, time series *prediction*, image *segmentation*, and *evaluation* through the estimation of plant descriptors, culminating in a validation process **(Figure 2 and Supplementary Figure S1).** Initially, we preprocessed all datasets to standardise image size, enhancing the training and predictive accuracy of our ML models. This standardisation is crucial for analysing small targets like early-stage Arabidopsis or minor leaf lesions formed in response to the pathogen.

### 2.2 Machine learning models for different modules of the AMULET pipeline

The processed images are analysed using a Faster Region-based Convolutional Neural Network (R-CNN) model^27^ to ensure accurate object *detection* **(Figure 2 and Supplementary Figure S1).** This model improves traditional structures by incorporating modifications to the feature extractor, anchor ratios, region of interest pooling, and loss functions, specially tailored to detect the specific characteristics of plant disease and phenotype traits effectively^28^. Using over 32,000 Arabidopsis plants, our ML model has been trained to ensure precise detection across all types of multi-well plates **(Supplementary Table S1)**. A universal automatic and accurate detection for all multi-well plates is crucial to ensure each plant is correctly identified and prepared for subsequent analysis.

To maintain positional consistency across different time points and plate distributions, we have developed a robust validation system that assigns a unique identifier to each plant (For more details, see **Supplementary Figure S2** and **Subsection-Data preprocessing**). This mechanism allows for precise tracking of individual plants over time, linking them with a database of previous measurements [**Supplementary material-Algorithm 1 ("Validation of Plant Positions")].**

Fast R-CNN excels in object detection and is computationally efficient^29,30^, essential for accurately identifying small Arabidopsis plants. While the Faster R-CNN model is widely used for disease identification across various plant species^31^, it has limitations in detecting small objects. To address this, we supplemented it with other models to identify efficiently, including three-day-old Arabidopsis seedlings (for further details, see **Subsection-Plant detection**).

The sequential imagery feeds into an image *prediction* model that forecasts the future appearance of each plant based on historical data (Figure 2). This predictive capability is essential for understanding plant growth patterns and potential future states. To achieve this, we integrated more than 3,000 Arabidopsis seedlings imaged at six consecutive time points into the model. The images were obtained from a modified HTP screening method based on De Diego et al.^8^ to study plant-pathogen interaction with the pathosystem model *Arabidopsis thaliana-Pseudomonas syringae (Pst)*. Our innovative non-invasive screening approach involved dropping bacterial suspension onto the leaves of two-week-old Arabidopsis, allowing a natural infection process (without infiltration) in which bacteria grow epiphytically (they can live and multiply on the outside of aerial surfaces of plants) and enter the plant through the stomata^32^. Images were automatically captured before infection (t_0_) and at intervals from 3 to 24 h post-inoculation (hpi) (from t_1_ to t_6_).

We selected the Simpler yet Better Video Prediction (SimVP)^33^ model for plant *prediction* because it is skilled at handling temporal image sequences. This model was compared against other leading models such as TAU^34^, ConvLSTM^35^, and PredRNNv2^36,37^ to ensure its suitability for our data temporal complexity (for further details, see **Supplementary Table S3**). Remarkably, SimVP predicted the appearance of infection symptoms two-time points ahead, detecting well infections at 9 hpi before they became visible to the naked eye at 24 hours (**Figure 2** and **Supplementary Figure S1)**. This model is rarely used in Natural Sciences^38^ and even less in Plant Science. It was only recently used to predict disease in *Ginkgo biloba* leaves at more advanced developmental stages when the damage in the tissue was already evident^39^. We demonstrated that the SimPV model was able to predict the progression of the leaves of young Arabidopsis seedlings infected with *Pst* inoculation 15 h before the lesions were visible, corroborating it as a very robust model suitable for retraining to predict changes in plant growth of other plant species.

Following prediction, the images underwent *segmentation* to isolate Arabidopsis rosette from the background, ensuring that subsequent analyses focus solely on plant evaluation without interference from background noise. We selected the pre-trained DeepLabV3+^40^ model integrated with an EfficientNet-B7 encoder after extensive testing with alternative encoders such as ResNet50 and ResNet101 **(Figure 2 and Supplementary Figure S1).** The EfficientNet-B7 stood out for its superior performance, ensuring precise segmentation for subsequent analysis phases ^41,42^. Due to the small size of both Arabidopsis seedlings and the lesions induced by *Pst* in this very early stage of colonisation, selecting this method proved to be crucial, as it enabled precise image segmentation, especially in our setup for accurately evaluating the image to define the organ or plant tissue that will be used for further analysis (**Supplementary Figure S1**). This task model, trained on a diverse dataset comprising over 4,000 images of Arabidopsis plants grown in well-plates, offered invaluable depth of pattern recognition. It avoided the recognition of artefacts reaching a precise plant segmentation, including for small objects **(Supplementary Figure S1)**. The DeepLabV3+ model has demonstrated strong performance, corroborated by its success in detecting plants and identifying various diseases across multiple plant species^43–46^. The EfficientNet encoders have also performed well in disease classification^47^. However, to this date, only one study has paired the DeepLabV3+ with the EfficientNet encoder as a part of the more complex pipeline that includes plant prediction^48^.

We use Dice loss and the Intersection over Union (IoU) score in the *segmentation* phase to evaluate performance. Dice loss measures the similarity between the predicted segmentation and the ground truth, complemented by the IoU score, which measures the overlap between the predicted and actual segmentation areas. This dual-metric approach enabled a comprehensive assessment of the model’s segmentation accuracy. Our training data set achieved exemplary performance: Dice loss of 0.0104 alongside an IoU score of 0.9948. Moreover, the evaluation of the validation set revealed a Dice loss of 0.0135 and an IoU score of 0.9932, and analysis of the test set showed a Dice loss of 0.0127 and an IoU score of 0.9936. Such results underscore the model’s robust and consistent performance, irrespective of the dataset being utilised. This uniformity in high-quality performance accentuates the model reliability in various contexts.

### 2.3 Statistical analysis of the plant descriptors

The final stage of our study involved a plant *evaluation* model trained to evaluate 24 distinct descriptors of plant morphology and colour changes derived from simple RGB images, analysed by the Data Analyzer (PSI, Czech Republic) software in our PlantScreen^TM^ systems. These traits were chosen due to the cost-effectiveness of RGB cameras and the heritability and reproducibility of morphological traits related to biomass, making them highly comparable between laboratory and field settings^49^. We statistically analysed these descriptors to understand the population variability of Arabidopsis seedlings (ecotype Col-0) responding to *Pst* inoculation at 24 hpi **(Figure 3).** Principal Component Analysis (PCA) captured significant variability in infection progression, with the two first principal components accounting for 47.7 % of the total model variance, distinguishing between plants with larger lesions (major infection) and smaller lesions (minor infection) **(Figure 3A).** This result underscored the need to increase the number of biological replicates in plant science studies to capture diversity.

**Figure 3.**
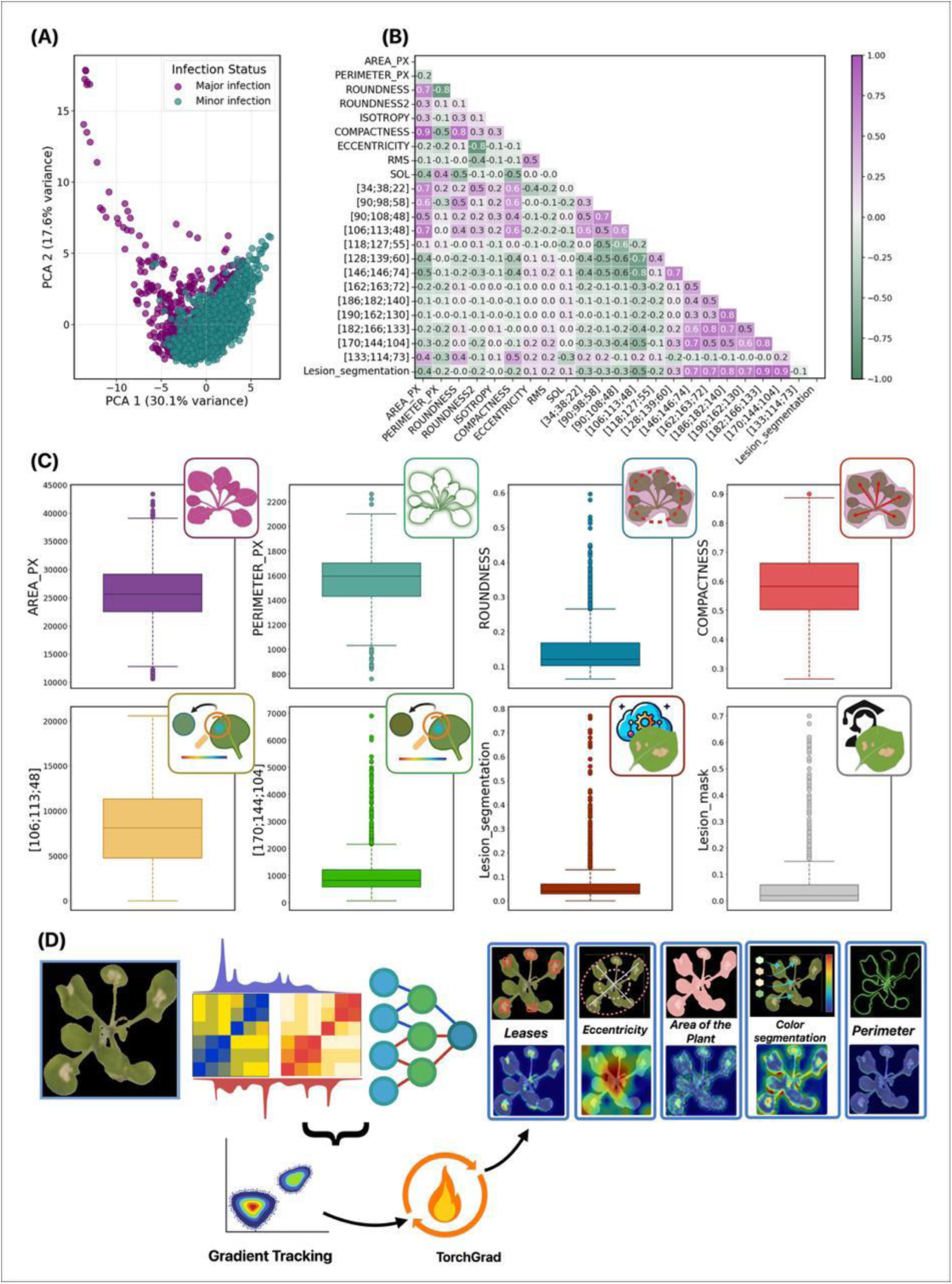
Statistical analysis and TorchGrad. (A) Principal component analysis (PCA) representing the population variability of two-week-old *Arabidopsis thaliana* Col-0 accession at 24 hpi with *Pseudomonas syringae* pv. *tomato* DC3000. (B) Correlation matrix between the morphological and colour-related descriptors obtained from the analysis of the RGB images. (C) Box plot of the most interesting descriptors. (D) TorchGrad highlights the most important image regions using the target gradients and their equivalence to the descriptors.

Correlation analysis of the 24 descriptors revealed two distinct groups **(Figure 3B).** The first group showed a positive correlation between traits like area (AREA_PX), roundness, compactness, and shades of green (e.g., 106: 113: 48 and 118: 127: 55) **(Figure 3B)**. The second group showed a positive correlation between lesion size (Lesion_segmentation) in Arabidopsis leaves at 24 hours post-*Pst* inoculation and a range of colours from brown to yellow **(Figure 3B)**. Notably, the perimeter and area were uncorrelated, highlighting that the perimeter provided unique and valuable information for inclusion in the model **(Figure 3B).** Perimeter and compactness have been reported to change rapidly in Arabidopsis plants under abiotic stress^50^. Similarly, our results indicated these parameters as promising markers for studying Arabidopsis responses to biotic stress, particularly in the model pathosystem *Arabidopsis thaliana-Pseudomonas syringae.* Overall, traits like rosette area, perimeter, roundness, specific colours, and lesion size give complementary insights into the plant response to *Pst* infection.

Further evidence of population variability in Arabidopsis response to *Pst* inoculation was observed in the box plot representations **(Figure 3C).** These visualisations showed the most variated distributions for descriptors such as area and perimeter (defined as the number of green pixels), roundness, compactness, and specific shades of green (RGB values 106: 113: 48, and 118: 127: 55). The box plots also indicated a normal distribution of the plant changes represented by these descriptors with only a few outliers **(Figure 3C).** Interestingly, strong positive correlations were found among the changes in the data obtained from several descriptors, such as area, compactness, and specific green colours (e.g., 106: 113: 48). This behaviour suggests a complex interaction between plant morphological parameters and colour traits in response to pathogen exposure.

Given the representativeness of the selected descriptors in our experimental setup, we used them in our plant *evaluation* models, which included more than 2,000 manually labelled plants. We fine-tuned various models for descriptor estimation, replacing the last classification layer with a regression layer. Using Microsoft ResNet50 as the teacher model and GhostNet v2 as the student model, we achieved high performance (R^2^ score of 0.9289), demonstrating the effectiveness of knowledge distillation in transferring robust feature detection capabilities to a more computationally efficient model. To ensure the robustness and efficiency of our pipeline, we employed Explainable AI (XAI) techniques, including TorchGrad and Gradient-weighted Class Activation Mapping (Grad-CAM). This technique highlighted important image regions using the target gradients to confirm that the plant evaluation model accurately focuses on relevant descriptors (**Figure 3D**). The comparison between Grad-CAM outputs and the selected descriptors confirmed that our ML model could effectively integrate the pertinent descriptors defining the Arabidopsis response to *Pst* inoculation. These techniques also facilitated a detailed evaluation of the model’s ability to identify and classify infected plants accurately, underscoring the potential of AMULET for rapid adaptation to different plant species and growth conditions^51,52^.

### 2.4. AMULET validation. A case study using in vitro potato plants

Although AMULET was initially developed using *Arabidopsis thaliana* for training and testing, the pipeline is specially designed for rapid adaptation to other plant species. The pre-trained models within the pipeline serve as robust backbones readily applicable across various contexts, from plant identification to evaluation. While these models offer a strong starting point, they typically require minor fine-tuning on small species-specific datasets to achieve optimal performances. This flexibility makes AMULET a highly versatile tool for agricultural applications, enabling a smooth transition between different species with minimal adjustments.

To demonstrate the versatility of AMULET, we adapted the plant *detection* and *prediction* tasks to analyse the growth of *in vitro* potato plants. *In vitro* techniques are crucial in potato research because they are the most commonly used methods for conserving global varieties in gene banks^53^ (https://cipotato.org/genebankcip/process/active_collection/). These techniques allow for easy, space-efficient propagation and maintenance of many plant species, from annuals to perennials^54^ (https://www.genebanks.org/the-platform/conservation-module/in-vitro/). With this aim, we developed a straightforward *in vitro* protocol for phenotyping *in vitro* potato plants using an affordable in-house small phenotyper equipped with Raspberry Pi Zero paired with a Raspberry Pi Camera V2 and LED lights for homogeneous illumination of the plants during image capture **(Supplementary Figure S4).** *In vitro* potato plants were grown in square assay tubes, with covers allowing gas exchange for more natural growth and avoiding condensation **(Supplementary Figure S4).** Potato plant development initiates from a single explant segment with a unique axillar bud, growing vertically into a new shoot with clear apical dominance. The explants were imaged vertically using a side-view RGB camera, presenting significant differences from our previous datasets **(Figure 4A).** Automating plant *detection* and *predicting* explant development under varying conditions can expedite the selection of optimal protocols. Moreover, integrating these techniques into the protocol for *in vitro* potato growth (or other relevant plant species conserved using *in vitro* approaches, including trees) facilitates the development of ultra-rapid screening methods to study potato resistance to different stressors using HTP screening.

**Figure 4.**
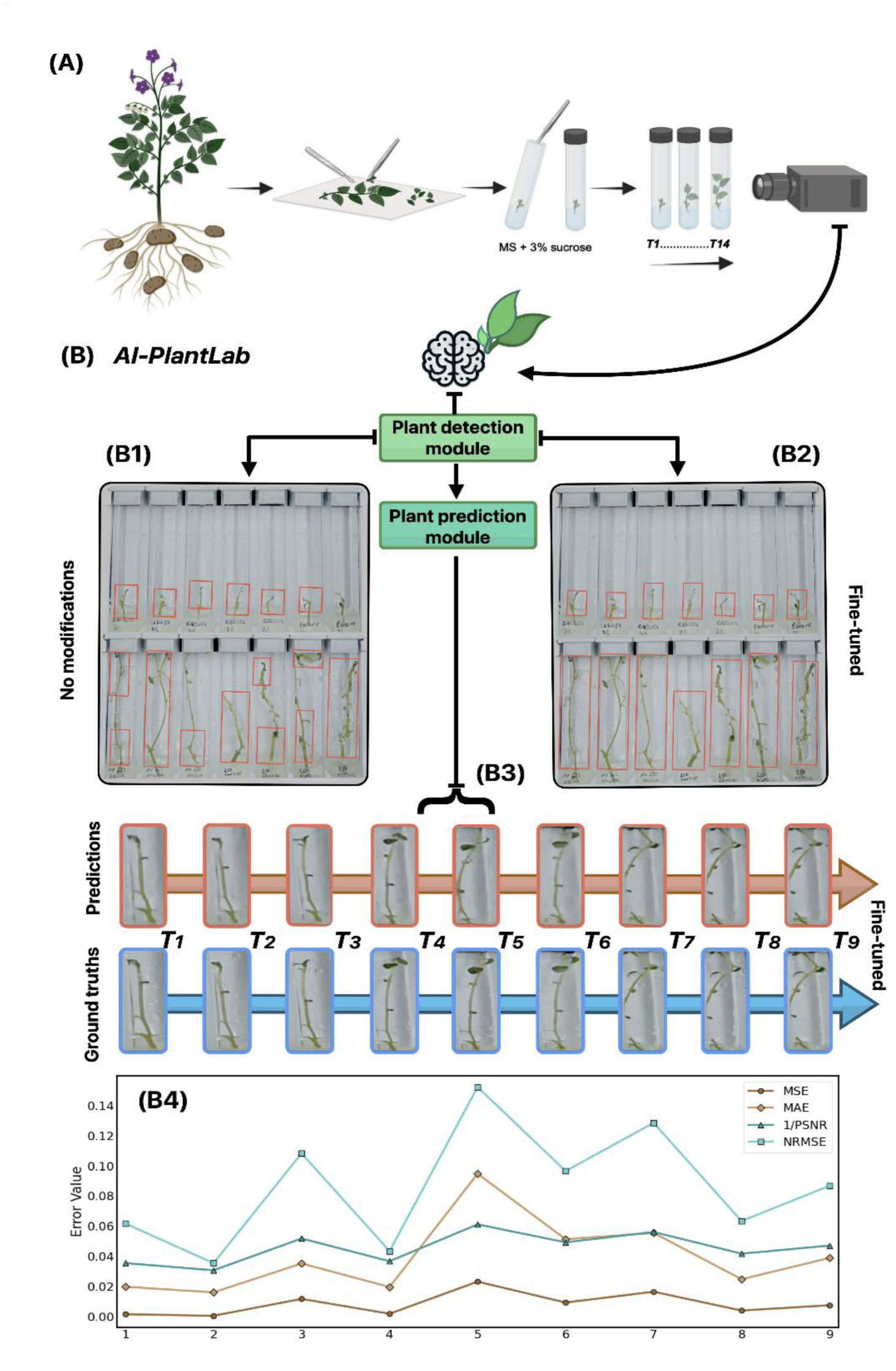
AMULET validation using *in vitro* potato plants. Adaptation of the AMULET pipeline to *in vitro* potato plants. (A) New protocol for *in vitro* potato imaging using side-view RGB camera. (B) Plant detection using the pre-trained model without modifications. (C) Improved plant detection after fine-tuning the model. (D) Fine-tuned image predictions for vitro potato plants: the blue section represents ground truth data from time steps *T1* to *T9*, and the red section illustrates the predictions made by the fine-tuned model. (D) Error metrics (MSE, MAE, 1/PSNR, NRMSE) for the predicted and ground truth images at each time step showcasing the impact of fine-tuning on the pre-trained AMULET modules.

Our study used approximately 100 pictures of potato plants grown *in vitro* over 15 days. We leveraged the pre-trained backbones from the plant detection and prediction phases. **Figure (4B1)** illustrates the initial detection performance of the *in vitro* potato plants capable of recognising individual plants. Although some mistakes were encountered, such as misidentifying separated parts of a plant or truncating the plant image, we addressed these issues by fine-tuning the pre-trained model on a smaller, targeted dataset of potato plants (100 annotated images for training and 40 for validation), allowing it to adjust parameters for improved accuracy **(Figure 4B2).** Leveraging the generalisation capability from extensive initial training, the model requires only a limited dataset, making the fine-tuning process efficient and effective, with solid performance gains achieved in minimal training time (hours).

Similarly, as illustrated in **Figure 4B3**, the image *prediction* phase for *in vitro* potato plants was fine-tuned. The blue section represents ground truth data from time steps *T*_1_ to *T*_9_, while the red section displays the predictions made by the fine-tuned model. **Figure 4B4** presents a comparison of several error metrics, namely Mean Squared Error (MSE), Mean Absolute Error (MAE), 1/Peak Signal-to-Noise Ratio (1/PSNR), and Normalised Root Mean Squared Error (NRMSE), quantifying the difference between the predicted plant images and the corresponding ground truths at each time-step. These metrics highlight the effectiveness of fine-tuning the pre-trained AMULET module in improving prediction capabilities. This adaptation underscores the efficiency and effectiveness of fine-tuning, particularly for tasks specific to a particular domain. AMULET has proved its adaptability to various plant species and growth conditions, providing a well-developed starting point for rapid fine-tuning. Moreover, the comprehensive nature of AMULET includes all necessary steps for processing plant phenotyping data, from detection to evaluation, streamlining the workflow, and saving valuable time once the imaging process concludes.

## 3. Conclusion

AMULET is a modular solution that significantly enhances the efficiency and speed of image processing from non-invasive phenotyping obtained using a low-cost RGB camera. It also incorporates prediction capabilities that offer multiple benefits for the stakeholders, ranging from the plant science community to crop growers, including shortened experiment timelines, early stress detection, and identification of unhealthy plants, allowing quicker treatment of stressed plants.

Trained on a large dataset featuring Arabidopsis at various developmental stages, including diminutive 3-day-old seedlings (2-4 mm diameter), AMULET has demonstrated remarkable adaptability across different plant species. The horizontal growth pattern of Arabidopsis simplifies monitoring, which contrasts with the vertical growth of *in vitro* potato plants used for validating AMULET. Despite initial challenges for precise detection of the potato explant, fine-tuning the model with only a minimal dataset achieved high accuracy, requiring far fewer resources and less time than building a new model from scratch, especially when the backbone is trained on a relevant, robust dataset. In this regard, AMULET used big data from a costly and sophisticated phenotyping system (PlantScreenTM XYZ) with a high-resolution RGB camera that is not widely accessible within the plant science community. However, AMULET’s pre-trained ML models are robust backbones that can be effectively applicable using a more accessible, low-cost phenotyper equipped only with an affordable RGB camera (Supplementary Figure S4).

AMULET has proven sensitive and effective in predicting the onset of diseases in Arabidopsis seedlings hours before visible symptoms appear. Integrating disease appearance prediction in agricultural management could permit preemptive interventions to improve plant health and minimise productivity losses. It also predicted the growth patterns of *in vitro* Arabidopsis and potato plants despite their different growth habits and contrasting plant body architecture. This capability holds significant promise for optimising protocols needed in Gene Banks, potentially accelerating the conservation and maintenance of a wide range of plant species, including recalcitrant ones.

The success of AMULET needs to be further challenged with a higher diversity of the scenarios of the infinite combinations of environmental conditions in which various plants and crops are growing. However, the proven AMULET’s ability to streamline the phenotyping process, from detection to data analysis, positions it as a transformative and breakthrough tool in plant science. Its versatility and efficiency could lead to advancements in breeding programs and agricultural management, empowering researchers and growers to implement proactive strategies that ensure plant vitality and yield optimisation, integrating and reducing effort and time post-plant imaging.

## 4. Material and Methods

### 4.1. Plant material and growth conditions

Seeds of *Arabidopsis thaliana* ecotype Col-0 were acquired from the Salk Institute Genomic Analysis Laboratory (http://www.signal.salk.edu/cgi-bin/tdnaexpress). Firstly, the seeds were surface sterilised and seeded on square plates (12 cm × 12 cm) containing 0.5× Murashige and Skoog (MS) medium (Murashige and Skoog, 1962) (pH 5.7) supplemented with a gelling agent 0.6% Phytagel (Sigma–Aldrich, Germany) and maintained for three days at 4 °C in the dark^8^. Afterward, the plates were transferred to a growth chamber with controlled conditions (22 °C, 16/8 h light/dark cycle, a light intensity of 120 µmol photons of PAR m^−2^s^−1^) and placed vertically. Four days later, seedlings of similar size were transferred under sterile conditions to 6- to 24-well plates with one seedling per well, and the plates were sealed with perforated transparent foil, allowing gas and water exchange. Each well contained different amounts of full MS medium (pH 5.7; supplemented with 0.6% Phytagel), according to the plate format. The plates were transferred to the Olophen platform equipped with the PlantScreen^TM^ XYZ system^8^. Arabidopsis seedlings were grown for a period ranging from seven to fourteen days, varying based on the number of wells in the plates, under controlled conditions (22/20 °C, 16/8 h light/dark cycle, a light intensity of 120 µmol photons of PAR m^−2^s^−1^, 60% relative humidity) on the Olophen platform.

We used six different potato varieties (Agria, Baraka, Carlita, Kennebec, Nu 633 Frisia, and Red Pontiac) multiplied *in vitro* for the case study. The material was obtained from healthy, disease-free mother plants grown *in vitro* and provided by NEIKER (Vitoria-Gasteiz, Spain). 7-10 mm mono-nodal segments containing one axillar bud each were cut from the mother plants, and leaves were excised. The nodes were then transferred to 7.7 × 7.7 × 9.7 cm magenta boxes (Sigma-Aldrich, Darmstadt, Germany) filled with 75 mL of full MS media, supplemented with 30 g/L sucrose, 8 g/L agar, pH = 5.7). Between 9 and 12 nodes were transferred into the magenta box to keep the plant material. Explants were kept in a Conviron CMP 6010 growth chamber (Conviron, Winnipeg, Canada) at 22 °C, under a light intensity of 80 µmol photons m^-2^ s^-1^, in long-day photoperiod (16 h day / 8 h night), until they had generated enough internodes (4 -5 weeks) for multiplying the explants to start the non-invasive phenotyping.

Plastic tubes (1.35 × 1.35 × 8.6 cm) (Tintometer GmbH, Dortmund, Germany) were filled with 4.3 mL of the same full MS described above for plant phenotyping. The air-tight caps were drilled four times each with a 2.5 mm bore to allow gas exchange. Each plant was subdivided into a 7-10 mm-long mono-nodal section containing one leaf and axillar bud. The two most basal and apical nodes were discarded. The leaves of the remaining nodes were excised, and one node was carefully inserted in the centre of each phenotyping tube until approximately a third of the node remained submerged in the media. The tubes were then closed with the perforated caps, and the gap on the top of each cap was filled with autoclaved glass wool (VWR International s. r. o., Stříbrná Skalice, Czechia) to minimise contamination. The tubes were kept for weeks in the growth chamber under the same conditions described above and photographed once per day using a Raspberry Pi Camera V2 and two stripes of cool-white LED lights for homogenous illumination. For imaging, the tubes were then placed in custom 3D-printed holders, designed with a curvature that allows the clear side of each tube to align perpendicular to the RGB camera (Supplementary Fig. S3B, C), thus minimising the plant area that gets occluded by the edge of the tube. These holders fit into the slot of a custom 3D-printed stand (Supplementary Fig. S3B, C), which can be placed in a fixed position in the phenotyper (Supplementary Fig. S3D). This system allows for obtaining homogeneous pictures in terms of lighting and distance from the camera.

### 4.2. *Infection* assay

Two-week-old Arabidopsis seedlings grown in 6 well-plates were used for the infection study. Some of the seedlings were inoculated with *Pseudomonas syringae* pv. *tomato* DC3000 *(Pst)* strain as described by Jasso-Robles et al.^55^, while the rest served as controls (non-inoculated plants). The bacterium was cultured in King’s B (KB) medium^56^ supplemented with 50 µg/mL rifampicin for 24 h at 28 °C. *Pst* was resuspended in 10 mM MgCl_2_ (pH 7.0) and adjusted to an OD600 of 0.2. Five µL of the bacterial suspension were inoculated on the true leaves of Arabidopsis seedlings. 5 µL of MgCl_2_ buffer was also applied to the remaining half of the Arabidopsis seedlings as a control.

### 4.3. Dataset Characteristics and Volume

All Arabidopsis images were captured using high-resolution RGB cameras integrated into our PlantScreen^TM^ XYZ or Compact system (Olophen Platform). For the initial phase of model training (plant *detection*), images of Arabidopsis seedlings grown in 6 to 24 well-plates at various developmental stages captured using high-resolution RGB cameras of our PlantScreen^TM^ XYZ system were used. The acquired images were initially annotated to identify key features, followed by extracting multiple descriptors, including morphological parameters such as the area (AREA PX) and perimeter (PERIMETER PX) calculated from binary images by counting pixels within the rosette and along its edge, respectively. Metrics like the relative growth rate, absolute growth rate, and colour-related indices (Green Leaf Index - GLI, Normalised Green-Red Difference Index - NGRDI, and Visible Atmospherically Resistant Index - VARI) were also calculated for each RGB image, as described by Ugena et al.^9^. These descriptors, encompassing both morphological traits and colour segmentation, were meticulously correlated with their respective images to facilitate accurate plant detection and analysis. 32,526 plants were used for the plant detection task **(Supplementary Table S1).**

For the *predictive* model task, a novel dataset comprising images of Arabidopsis seedlings grown in 6-well plates. Two-week-old plants inoculated with *Pst* or MgCl_2_ as control were photographed at predetermined intervals before and after inoculation (*t*_0_ to *t*_5_; 0, 3, 6, 9, 12, and 24 hours) using a high-resolution RGB camera installed in our PlantScreen^TM^ Compact system to include a comprehensive range of conditions encompassing infected and healthy plants.

Expert verification was conducted on a separate test dataset, focusing on the morphology and health status of *Arabidopsis thaliana* (3,312 plants) to validate the predictive accuracy of the models **(Supplementary Table S1).** In this case, two additional features related to the size of the leaf lesions were also included for higher accuracy.

For the *segmentation* task, additional 3,306 plants were carefully selected and manually segmented to capture the diversity related to the issue, comprising 2,298 infected and 1,008 non-infected Arabidopsis seedlings **(Supplementary Table S1)**. An additional 546 infected and 545 non-infected plants segmented by our model were also added to this dataset, and each image underwent rigorous verification to ensure relevance and accuracy.

Finally, for the plant *evaluation* task, a new dataset of 2,298 *Pst*-infected plants was chosen explicitly for annotation and further analysis **(Supplementary Table S1).** This dataset included additional morphological descriptors such as roundness, roundness2, and isotropy, which describe plant circularity and eccentricity, and RMS, which indicate symmetry^57^. Parameters like compactness and SOL, which depend on the distance of the edge to the centre, were also used. Furthermore, automated (Lesion segmentation) and manual (Lesion Mask) segmentation of each plant lesion area was included in the annotated descriptors for this phase for a more complex description of the plant health status.

### 4.4. AMULET Details

With proper data preprocessing, partitioning, and augmentations **(see Subsection- 4.5. Preprocessing and Augmentation),** we utilised the initial RGB imagery of shape (*N* × 3 × 1920 × 2560), where *N* is the number of gathered images to train the Faster Region-based Convolutional Neural Network (Faster-RCNN) for the plant *detection* task. The Faster-RCNN model, as depicted in **Supplementary Figure S1**, integrates two key components: a region proposal network (RPN) for generating region proposals and a deep convolutional neural network (CNN) for the classification and bounding box refinement of these proposals. The RPN directly utilises the feature maps obtained from the input images to predict object bounds and objectness scores at each position. The CNN then processes these proposals, classifying the objects within the proposed regions and refining their locations. During the training and testing process, the Box Regression, Classification, Objectness, and RPN Box Regression losses-scores were monitored and logged to ensure the robustness of the training process for exact loss values and other parameters obtained in the training of the pipeline **(Supplementary Table S1**).

For the pivotal task of plant *prediction*, which takes as input image time series of shape (*N* × *T* × 3 × 224 × 224), where *T* is the number of plant images from previous measurements, we selected the Simpler yet Better Video Prediction model (SimVP). The SimVP framework utilises a CNN-based approach comprising an Encoder, Translator, and Decoder. The Encoder extracts spatial features through *N*_*s*_, Conv, Norm, and ReLU blocks, operating on *C* channels across dimensions (*H*, *W*). The Translator captures temporal dynamics using *N*_*t*_ Inception modules, convolving *T* × *C* channels over (*H*, *W*). The Decoder reconstructs the future frames with Ns unConv Norm ReLU blocks, targeting *C* channels on (*H*, *W*). The simplified global structure of the model is available in **Supplementary Figure S1**.

For the training of the plant *segmentation* step, which adopts input data of shape (*N* × 3 × 224 × 224), the architecture of DeepLabV3+ was chosen integrated with the EfficientNet-B7 encoder. The simplified architecture of the DeepLabV3+ is depicted in **Supplementary Figure S1**. It integrates multi-scale contextual information through atrous convolution applied at various scales within the encoder module, facilitated by the EfficientNet-B7 advanced feature extraction capabilities. To highlight the effectiveness of this trained model, plant images were segmented using a conventional segmentation system, providing a baseline for comparison and displaying images segmented by our fine-tuned DeepLabV3+ model. See **Supplementary Table S2** for the obtained loss summary for each step.

For the final phase of the AMULET pipeline, we trained a plant *evaluation* model that takes as input segmented images (*N* × 3 × 224 × 224) and their corresponding targets in the form of 24 unique descriptors (*N* × 24). To estimate these crucial descriptors from images, we separately trained a set of models, namely Vision Transformer (ViT)^58^, CoAtNet^59^, ResNet50, and GhostNet^60^, from which the best performance was exhibited by the ResNet50 **(Supplementary Table S2**). Furthermore, we employed knowledge distillation between combinations of the models mentioned, which resulted in the best combination of ResNet50 as the teacher and GhostNet v2 as the student, significantly outperforming other variations **(Supplementary Figure S1 and Table S2**). Distillation knowledge is transferred from ResNet50 to GhostNetV2-160, utilising true and soft labels to guide the student model in replicating the teacher’s analytical prowess. This approach ensures that the student model achieves comparable accuracy in terms of computational efficiency. Validating outputs from the previous tasks is quite straightforward; however, in this case, the model outputs just a list of values guided by the *R*^2^ score.

### 4.5. Preproccesing and Augmentation

Data preprocessing is essential for preparing raw images and auxiliary inputs, significantly impacting algorithm performance in training and deployment. This is particularly important for complex data like the Arabidopsis dataset. The normalisation preprocessing step is applied for Plant Detection, Prediction, Segmentation, and Evaluation, ensuring a uniform range for pixel values. Additionally, we standardise descriptors extracted from segmented images, which serve as target labels in the Plant Evaluation step, to ensure balanced contributions to the model’s learning process. Furthermore, a plant position validation procedure assigning a unique identifier to each plant following a systematic convention was created as detailed in **Supplementary material- Algorithm 1** and illustrated in **Supplementary Figure S1**. This identification begins with the plant located at the top-left and progresses horizontally across the image.

Moreover, dynamic thresholding was used to accommodate plant growth between consecutive time points. This method involves evaluating the plant size bounding box in sequential images and making necessary adjustments based on a predefined growth rate threshold. **Supplementary material- Algorithm 1** details the procedural steps for detecting, organising, and assigning unique indices to plants within a given image. These steps adhere to a systematic workflow that includes loading the image, applying a pre-trained model, extracting and sorting detected plants, and assigning positions based on a vertical threshold and row-wise organisation. Additionally, **Supplementary Figure S2** demonstrates its practical application. The centring equation inside **Supplementary material- Algorithm 1** that calculates the centre of the bounding box is defined as follows:

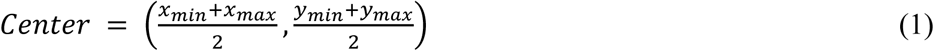

where *x*_*min*_ and *x*_*max*_ represent the minimum and maximum x-coordinate values of the bounding box, determining the leftmost and rightmost boundaries of the object detected in the image, respectively. Similarly, *y*_*min*_ and *y*_*max*_ denote the minimum and maximum y-coordinate values of the bounding box, defining the bottom and top boundaries of the object, respectively (**see Supplementary Figure S2 and Supplementary material- Algorithm 1**). Additionally, we examined the plant changes and development and excluded Arabidopsis plants with no or minimal growth in subsequent images from the dataset. This systematic approach ensures that our models remain sensitive to the natural variability in plant sizes and maintain the integrity of temporal data analysis.

Despite generating high-quality data, data augmentation remains essential for training models. Introducing diverse variations through augmentation significantly enriches the dataset, enhancing model performance. Augmentation strategies are applied at each pipeline stage, ensuring comprehensive enhancement of the dataset diversity. Several augmentations were used throughout the pipeline, such as brightness adjustments to simulate diurnal variations in sunlight exposure at different times of the day. Gaussian blur is applied to emulate the blurring effects that might occur during data collection, introducing the model to potential variances in image clarity.

Additionally, random positioning augmentation ensures geometric variability of plant placements within images, enriching the dataset with a broader range of positional contexts. Rotation augmentations are introduced at certain pipeline stages to account for orientation variability. For a summary of augmentations used in each step of the training pipeline, see **Supplementary Table S3**.

## Funding

This publication was funded by the project PATAFEST funded by the European Union under the grant number 101084284. The views and opinions expressed are solely those of the author(s) and do not necessarily reflect those of the European Union or the European Research Executive Agency (REA). Neither the European Union nor the granting authority can be held responsible for them. This publication was also funded by the project JG_2024_036, implemented within the Palacky University Young Researcher Grant.

### Acknowledgements

We thank Jana Nosková and Andrea Hybenová for their technical support. We thank NEIKER, particularly Dr. Jose Ignacio Ruiz de Galarreta and Dr. Amaia Ortiz-Barredo, for providing the *in vitro* potato material.

## Author contribution

J.Z., F.G., L.S., V.S., and N.D.D. conceived the manuscript’s structure. J.Z and N.D.D. led the writing of the manuscript and experimental work. P.K., F.I.J-R. and I.S-F. contributed to experimental work regarding plant phenotyping. J.Z and L.K. contributed to the experimental work regarding machine learning models. J.Z., L.K., P.K., F.G., L.S., V.S., and N.D.D. contributed to the writing of the manuscript. All authors reviewed and agreed on the manuscript before submission.

## Data and materials availability

The datasets generated and/or analysed during the current study are available from the corresponding author upon reasonable request.

