## Supplementary Material for "Decrypting the complex phenotyping traits of plants by machine learning"

Methodology article- Supplementary Material

### Supplementary Tables

**Supplementary Table S1** Summary of plant data across detection, prediction, segmentation, and evaluation stages. Detection includes image counts, plate types, foil presence, and estimated plant numbers for infected and non-infected conditions. Prediction provides a time-step analysis of plant estimates. Segmentation compares manual and model-based labelling for accuracy verification. Evaluation details were manually labelled, and plant counts were verified with statistical references (see Figure 3).

**Plant Detection**

| Status | Image Count | Plate Type | Foil Presence | Est. num. of plants |
|---|---|---|---|---|
| Infected | 387 | 6 | ✕ | 2322 |
| Infected | 235 | 12 | ✔ | 2820 |
| infected | 259 | 24 | ✔ | 6216 |
| Non-infected | 42 | 24 | ✕ | 1008 |
| Non-infected | 840 | 24 | ✔ | 20160 |
| Estimated plant $\sum 32{,}526$ | | | | |

**Plant Prediciton**

| Status | Image Count | Plate Type | Time Step [h] | Est. num. of plants |
|---|---|---|---|---|
| Infected | 32 | 6 | 0 | 192 |
| Infected | 32 | 6 | 3 | 192 |
| Infected | 32 | 6 | 6 | 192 |
| Infected | 32 | 6 | 9 | 192 |
| Infected | 32 | 6 | 12 | 192 |
| Infected | 32 | 6 | 24 | 192 |
| Non-infected | 60 | 6 | 0 | 360 |
| Non-infected | 60 | 6 | 3 | 360 |
| Non-infected | 60 | 6 | 6 | 360 |

| | | | | |
|---|---|---|---|---|
| Non-infected | 60 | 6 | 9 | 360 |
| Non-infected | 60 | 6 | 12 | 360 |
| Non-infected | 60 | 6 | 24 | 360 |
| Estimated plant ∑ 3,312 | | | | |

**Plant Segmentation**

| Status | Num. of Plants | Manually labeled | Labeled by model | Verified |
|---|---|---|---|---|
| Infected | 2298 | ✔ | ✕ | ✔ |
| Non-infected | 1008 | ✔ | ✕ | ✔ |
| Infected | 546 | ✕ | ✔ | ✔ |
| Non-infected | 545 | ✕ | ✔ | ✔ |
| Estimated plant ∑ 4397 | | | | |

**Plant Evaluation**

| Status | Num. of Plants | Manually labeled | Verified | Statistic |
|---|---|---|---|---|
| Infected | 2298 | ✔ | ✔ | See Figure 3 |

**Supplementary Table S2.** Performance metrics for plant detection, segmentation, prediction, and evaluation across various models and tasks. Metrics include Box Regression, Classification, Objectness, and RPN Box Regression for detection; Intersection over Union and Dice loss for segmentation; MSE for prediction with SimVP, TAU, ConvLSTM, and PredRNNv2; and $R^2$ values for evaluation using standalone and combined models like ResNet50, ViT, CoatNet, and GhostNet v2.

| **Plant Detection** | **Train** | **Test** |
|---|---|---|
| Box Regression | 0.130 | 0.141 |
| Classification | 0.058 | 0.058 |
| Objectness | 0.011 | 0.015 |
| RPN Box Regression | 0.028 | 0.029 |

| **Plant Segmentation** | **Train** | **Test** |
|---|---|---|
| Intersection over Union | 0.995 | 0.993 |
| Dice loss | 0.010 | 0.013 |

| **Plant Prediction** | **Train** | **Test** | **Test MSE** |
|---|---|---|---|
| SimVP | **0.0051** | **0.0470** | **191.00** |
| TAU | 0.0142 | 0.0499 | 200.97 |
| ConvLSTM | 0.0289 | 0.0742 | 328.06 |
| PredRNNv2 | 0.0146 | 0.0512 | 202.86 |

| **Plant Evaluation** | $R^2$ **test** | $R^2$ **Distillation test** |
|---|---|---|
| Microsoft ResNet50 (alone) | **0.912** | - |
| Google ViT (alone) | 0.894 | - |
| CoatNet (alone) | 0.820 | - |

| | | |
|---|---|---|
| GhostNet v2 (alone) | 0.896 | - |
| Microsoft ResNet50 + CoatNet | - | 0.874 |
| Microsoft ResNet50 + GhostNet v2 | - | **0.929** |
| Google ViT + CoatNet | - | 0.869 |
| Google ViT + GhostNet v2 | - | 0.897 |

**Supplementary Table S3.** Overview of data augmentation techniques customised for each processing block in Our Pipeline. Notation: ✓ corresponds to applied, × corresponds to not applied.

| Augmentation Technique | Detection | Prediction | Segmentation | Evaluation |
|---|---|---|---|---|
| Gaussian Blur | ✓ | ✓ | ✓ | ✓ |
| Brightness Contrast | ✓ | ✓ | ✓ | ✓ |
| Random Sharpen | ✓ | ✓ | ✓ | ✓ |
| Random Positioning | ✓ | × | × | × |
| Random Rotation | × | ✓ | ✓ | × |
| Gaussian Noise | × | × | ✓ | × |
| Vertical Flip | × | × | × | ✓ |
| Horizontal Flip | × | × | × | ✓ |

**Supplementary material- Algorithm 1**

**Algorithm 1:** Validation of Plant Positions

**Input:** N plate setup Image;

**Output:** assigned positions of the plants;

**1. Initialise: M** ←Load image and corresponding metadata ;

**2. Image pre-scan:** Apply the NN model to obtain (bounding boxes, labels, scores) ;

**3. Image Extraction:** ;

**for** *plant* ∈ ***M*** **do**

a. Calculate the *center* of the bounding box; see **Eq 1** ;

b. Match (bounding box, label, score) to *center* ;

**end**

**4. Sort and Organise: ;**

a. Plants in row-wise organized← Sort(plants, y-coordinate, centers) ;

b. Organise plants for rows and remove plants that do not match ;

**5. Assign Positions: ;**

a. assign indices to the plants obtained from the previous step ;

b. **P** ← pre-screen plants based on predetermined criteria (e.g., top num. expect.) ;

**6. Naming Convention: ;**

**For** *plant* ∈ ***P*** **do**

plant ← filename: {experiment name, image number,plant population, index} ;

**end**

### Supplementary Figures

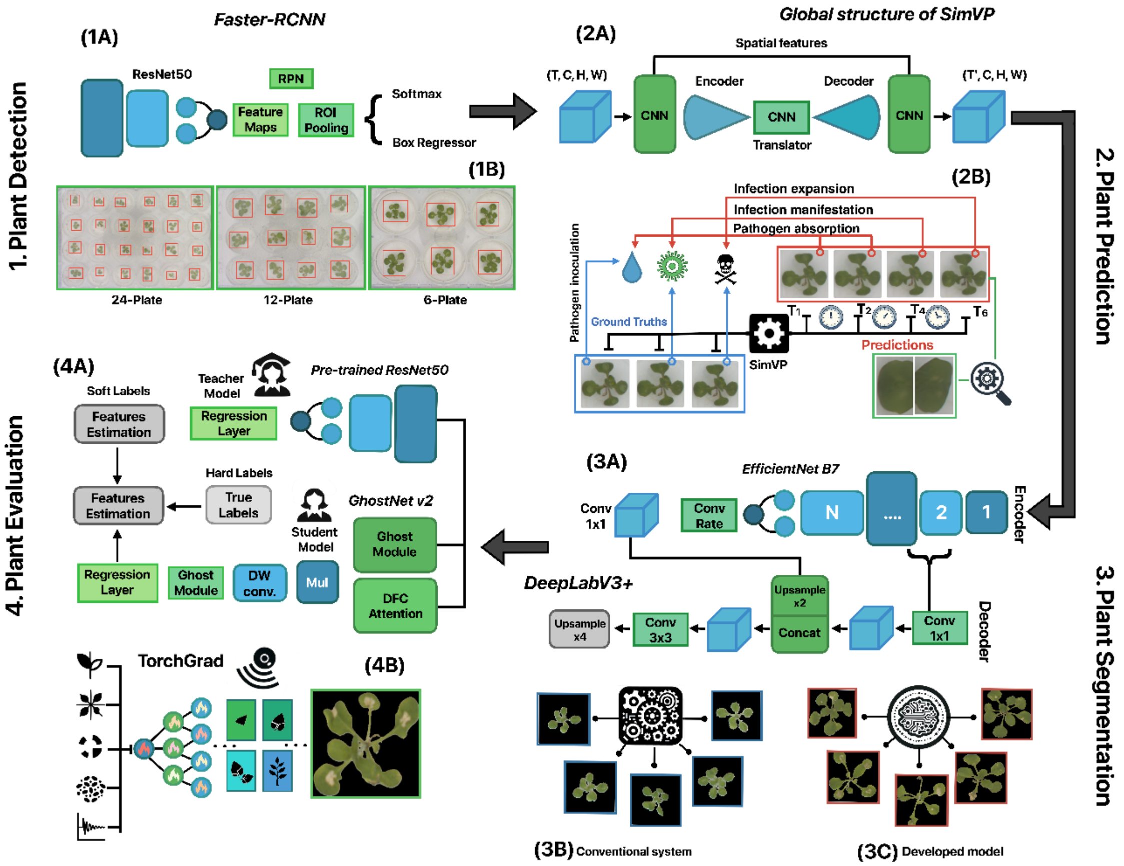


**Supplementary Figure S1**. Detailed AMULET pipeline. **(1) Plant *detection* task-** (1A) Faster-RCNN model architecture with Region Proposal Network (RPN) and deep CNN for classification and bounding box refinement and (1B) plant detections by Faster-RCNN grown into 6, 12, and 24-well plates, showing accurate identification and localisation. **(2) Plant *prediction* task-** (2A) SimVP model structure with Encoder, Translator, and Decoder components and (2B) SimVP predictions capturing the spread of infection over time, shown alongside ground truth images. **(3) Plant *segmentation* task-** (3A) Simplified DeepLabV3+ architecture with EfficientNet-B7 encoder using atrous convolution for multi-scale information, (3B) baseline plant images segmented with a conventional system, and (3C) plant images segmented by the fine-tuned DeepLabV3+ model showing improved accuracy. **(4) Plan evaluation-** (4A) Knowledge distillation from ResNet50 to GhostNetV2-160, illustrating analytical prowess transfer and (4B) TorchGrad visualisation highlighting model's focus on crucial features for estimating plant descriptors.

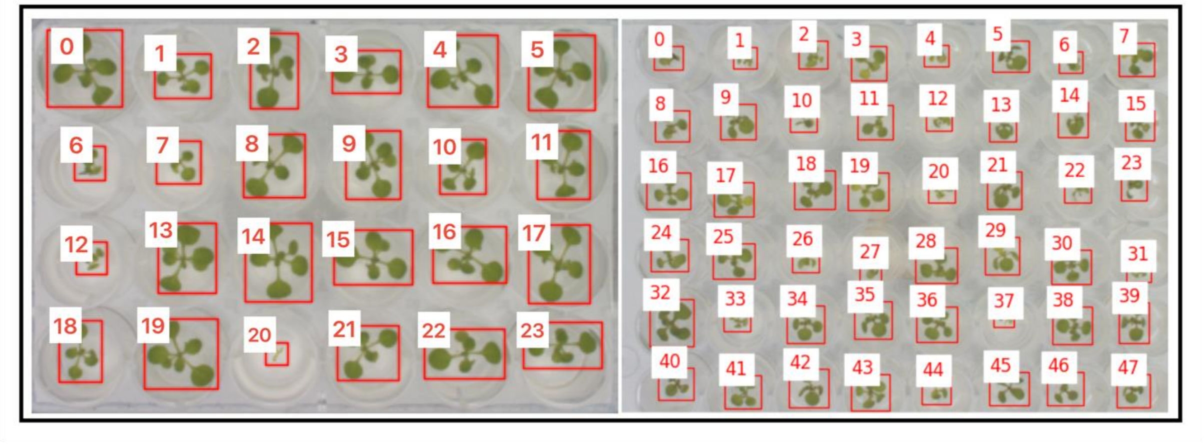


**Supplementary Figure S2.** Example of plant position assignment. It demonstrates the assigned indexing for plants, with the top image illustrating the indexing for a 48-well plate layout and the bottom image for a 24-well plate layout, showcasing the necessity of adaptability in the algorithm's design across different plate sizes.

.

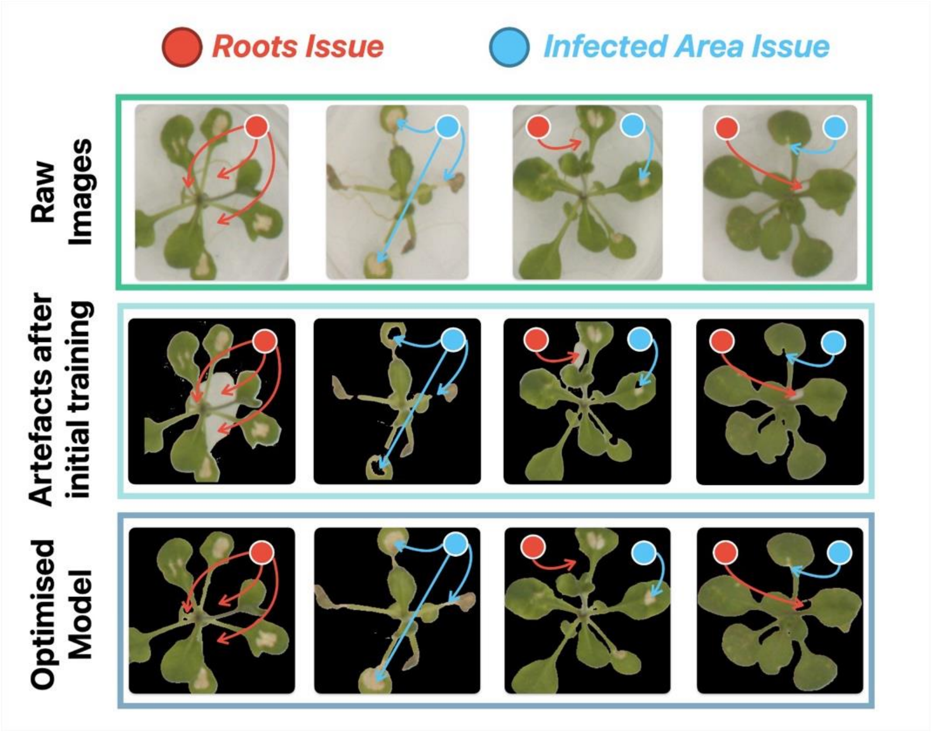


**Supplementary Figure S3**. Artifacts appearing during the plant *segmentation* task. The images showed the background effect and the complexity of correct plant segmentation.

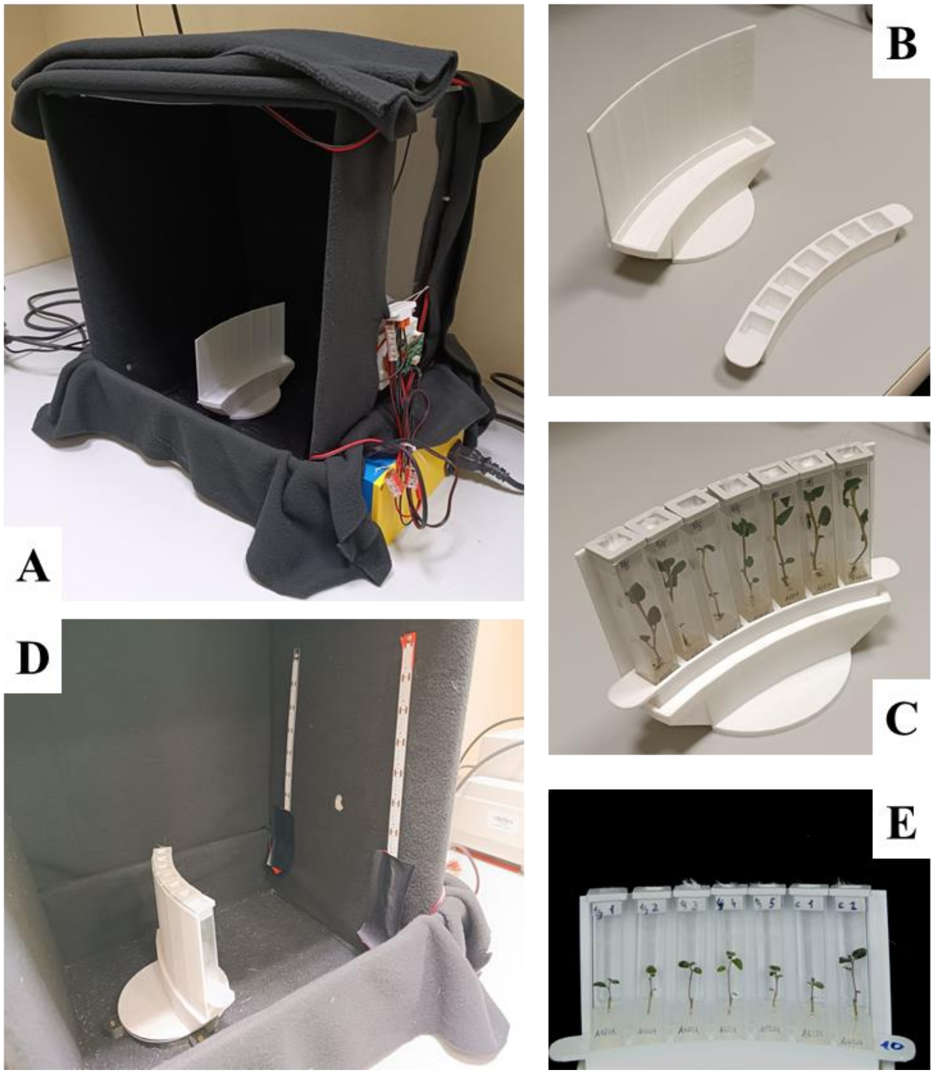


**Supplementary Figure S4.** (A) Custom phenotyper employed to obtain the potato images. (B) Custom 3D-printed stand and tube holder. (C) Example of *in vitro* plants placed on the homemade holder and stand. (D) Example of *in vitro* plants ready for imaging in front of the raspberry Pi Camera V2 and cold white LED stripes. (E) Picture of *in vitro*-grown potato plants taken using our custom phenotyping setup.
